# Knocking out two polyphenol oxidase genes significantly improves recombinant protein purification in *Nicotiana benthamiana*

**DOI:** 10.1101/2025.09.21.677675

**Authors:** Hai-Ping Diao, Hui-Xin Meng, Xue-Jiao Xu, Yong-Feng Guo, Inhwan Hwang, Shi-Jian Song

**Affiliations:** Qingdao Municipal Key Laboratory of Plant Molecular Pharming, Tobacco Research Institute, Chinese Academy of Agricultural Sciences, Qingdao, China; Beijing Life Science Academy (BLSA), Beijing, China; Department of Life Science, Pohang University of Science and Technology, Pohang, Korea

**Author notes:** These authors contributed equally to this work (Hai-Ping Diao, Hui-Xin Meng).

**Keywords:** Plant molecular pharming, *Nicotiana benthamiana*, Polyphenol oxidases, Protein aggregation, Protein purification

## Abstract

Efficient purification remains one of the major bottlenecks in the development of plant-based systems for recombinant protein production. The complex metabolites, particularly polyphenols, which usually cause recombinant protein aggregation during purification. In this study, we identified two key polyphenol oxidase genes, *PPOa* and *PPOb* from *N.benthamiana* as responsible for these effects. Using CRISPR/Cas9, we generated two *ppoa;ppob* double knockout lines that significantly improved the purification of functional proteins like SARS-CoV-2 Spike trimer and influenza HA trimer. These lines showed reduced polyphenol-protein interactions, minimized aggregation, and higher purification yields. Our work establishes a clean, high-efficiency *N. benthamiana* chassis for scalable recombinant protein production.

## Introduction

Chassis engineering plays a pivotal role in driving progress in synthetic biology. Advances in the design and optimization of microbial and mammalian cell-based chassis have enabled the efficient production of diverse bioproducts, including opiates(Nakagawa et al., 2016), vitamin B_12_(Fang et al., 2018), tropane alkaloids(Srinivasan and Smolke, 2019), antibodies(Chan et al., 2025), subunit vaccines(Tian et al., 2021; Xu et al., 2022; Weidenbacher et al., 2023). In contrast to microbial or animal cell chassis, plant-based chassis are inherently more complex due to the presence of diverse endogenous proteins and secondary metabolites, such as terpenoids and polyphenols. These intrinsic components present significant challenges for the purification of recombinant proteins from crude plant extracts. Studies have reported that purification can account for up to 80% of the total production cost for plant-derived recombinant proteins(Buyel, 2019).

Our previous researches revealed that plant-derived recombinant proteins, such as influenza virus HA trimers (HA_t_)(Song et al., 2021) and SARS-CoV-2 Spike trimers (S_t_)(Song et al., 2022b), exhibit strong interactions with plant polyphenols during Ni^2+^-NTA affinity purification. These interactions lead to severe enzymatic browning and protein aggregation, which are resistant to removal by conventional chromatographic methods, thus substantially impairing purification efficiency and compromising product quality. The interactions between polyphenols and proteins are well recognized and extensively studied in the field of Food Science, occurring through both covalent (irreversible) and non-covalent (reversible) mechanisms(Lu et al., 2025). Covalent binding typically involves the oxidation of polyphenols to ortho-quinones by polyphenol oxidases (PPOs) in the presence of oxygen. These quinones then react with free amino- or thiol-groups on proteins, resulting in irreversible protein– polyphenol complexes(Zhang et al., 2024). Building on this background, we raise the following scientific question: can targeted identification and knockout of polyphenol oxidase genes in *N.benthamiana* suppress the conversion of polyphenols into o-quinones, thereby preventing their interaction with recombinant proteins?

## Results

Previous studies identified six candidate PPO genes in *Nicotiana benthamiana* through genome annotation and transcriptome sequencing(Mahadevan et al., 2025). Transcriptome analysis of 4-week-old leaves revealed that *NbL17g16540*, which we named *PPOa* in this study, exhibited strong expression. Another three genes, *NbL10g18390(PPOb), NbL06g06640(PPOc), and NbL17g16550 (PPOd)* show high protein sequence similarity (Figure 1A), but minimal expressions were designated. To assess gene expressions across developmental stages, we extracted RNA from leaves at three different positions and performed qRT-PCR (Figure 1B and 1C). Transcript levels of *PPOs* were consistent high across leaf positions, but *PPOb, PPOc,* and *PPOd* consistently showed Ct values above 35 (Figure 1D), confirming very low expression level, in line with transcriptomic data. To investigate their functional roles, we first designed two 300-nucleotide fragments from *PPOa* for virus-induced gene silencing (VIGS) assay. The first construct, *TRV2::PPOa-1*, differs from the homologous region of *PPOb* by only 3 nucleotides and could theoretically target *PPOb* simultaneously. In contrast, the second construct, *TRV2::PPOa-2*, contains 9 dispersed differences in the homologous region, suggesting that its silencing effect on *PPOb* would likely be limited (Figure 1E, Figure S1). Both *TRV2::PPOa-1* and *TRV2::PPOa-2* plants showed no growth or developmental phenotypes compared to *TRV2::GUS* plants (Figure 1F). Notably, *TRV2::PPOa-1* rather than *TRV2::PPOa-2* led to a marked reduction in enzymatic browning and protein aggregation (Figure 1G and 1H), indicating a combined role of *PPOa* and *PPOb* in mediating these effects. These results guided the selection of targets for genes knockout.

**Figure 1.**
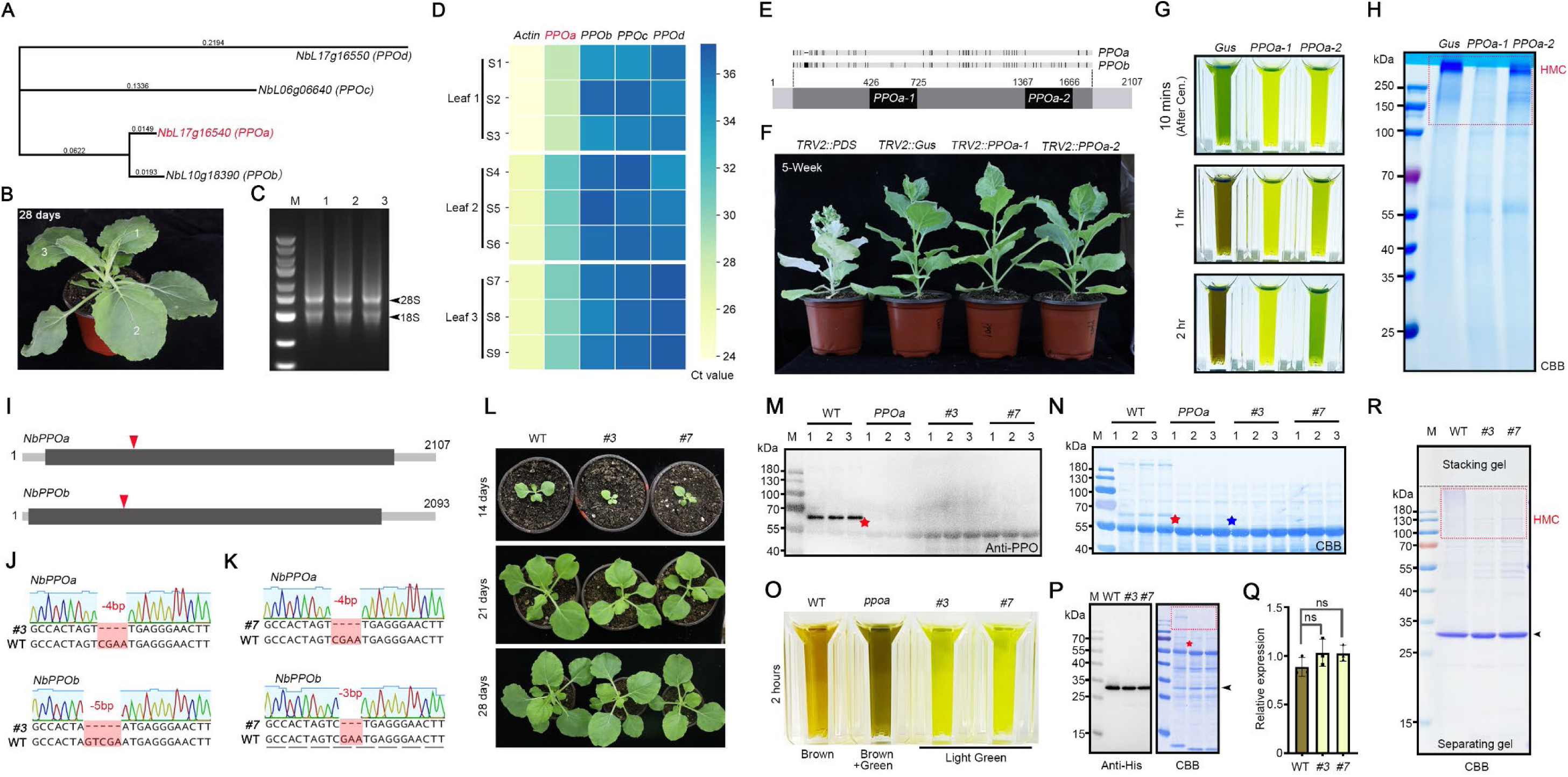
Identification of key *NbPPO* genes and generation of knockout mutants. **(A)** Phylogenetic tree of four *NbPPO* genes from the NbLab360 database. **(B)** 28-day-old *N. benthamiana* plants showing the three leaves collected for RNA extraction. **(C)** RNA quality assessment by agarose gel electrophoresis. **(D)** Ct values of the four PPO genes in different leaves of 28-day-old plants. **(E)** Schematic of the *NbPPOa* gene, highlighting two distinct regions selected for VIGS silencing. Light grey: introns; dark grey: exons; black: 300 bp gene fragments used to construct the TRV2 vectors. **(F)** Growth phenotypes of *TRV2::PDS, TRV2::Gus, TRV2::PPOa-1*, and *TRV2::PPOa-2* plants. **(G)** Total soluble extracts from 5-week-old *TRV2::Gus, TRV2::PPOa-1*, and *TRV2::PPOa-2* plants incubated at room temperature for 10 min, 1 h, and 2 h, photographed in cuvettes. **(H)** Total soluble extracts from *TRV2::Gus, TRV2::PPOa-1*, and *TRV2::PPOa-2* plants incubated with Ni²⁺-NTA beads for 15 min at 4 °C, followed by three washes. Beads were then boiled in 5× SDS loading buffer for SDS–PAGE and Coomassie Brilliant Blue (CBB) staining. HMC: high molecular complex. **(I)** Schematic of *NbPPOa* and *NbPPOb* genes, showing introns (light grey), exons (dark grey), and gRNA target sites (red arrows). **(J, K)** Mutation maps of two double-knockout lines, *ppoa;ppob#3* and *ppoa;ppob#7*. **(L)** Growth phenotypes of WT, *ppoa;ppob#3*, and *ppoa;ppob#7* plants. **(M, N)** Western blot and CBB staining analysis of total soluble extracts from WT, *ppoa*, *ppoa;ppob#3*, and *ppoa;ppob#7*. Red asterisks mark the endogenous PPOa band; blue asterisks indicate other endogenous PPOs, likely PPOb. **(O)** Total soluble extracts from WT, *ppoa*, *ppoa;ppob#3*, and *ppoa;ppob#7* plants after 2 h incubation at room temperature, photographed in cuvettes. **(P)** Western blot and CBB analysis of recombinant GFP expressed in WT and *ppoa;ppob*. Red asterisks mark endogenous PPOa; black arrow indicates recombinant GFP. The expression quantification result is shown in **(Q).** Statistical differences were determined by t-test. Error bars represent SEM. **(R)** CBB analysis of recombinant GFP purified from WT and *ppoa;ppob*. Red dashed box: high molecular complex; black arrow: recombinant GFP.

Building on the VIGS results, we developed a CRISPR-Cas9 construct to simultaneously knock out *PPOa* and *PPOb*. This yielded two independents *ppoa;ppob* double mutant lines (#3 and #7, Figure 1I - 1K) and several single *ppoa* mutants. Phenotypic analysis of T3-generation *ppoa;ppob* mutants showed delayed germination compared to WT plants. By the fourth week, however, their growth was comparable to WT, and no significant differences were observed in later reproductive development (Figure 1L). Western blot and CBB analysis confirmed that endogenous PPO proteins were undetectable in *ppoa;ppob* double mutants and weakly detectable in *ppoa* single mutants (Figure 1M and 1N). Correspondingly, enzymatic browning was pronounced in WT extracts, mild in *ppoa*, and absent in *ppoa;ppob* (Figure 1O). To assess the impact on recombinant protein purification, we expressed a GFP-His protein in both WT and *ppoa;ppob* plants. While expression levels were similar (Figure 1P and 1Q), Ni²⁺-NTA purification followed by SDS-PAGE and CBB analysis revealed significantly reduced protein aggregation in the *ppoa;ppob* plants which is marked as high-molecular complex (HMC) (Figure 1R). This indicates that eliminating *PPOa* and *PPOb* enhances recombinant protein quality by mitigating polyphenol-mediated aggregation.

Recombinant HA_t_ and S_t_ expressed in *N.benthamiana* frequently formed severe aggregates during purification. Although adding PVPP during liquid nitrogen grinding helped adsorb some endogenous polyphenols and modestly improved protein quality(Song et al., 2022b), this approach is impractical for large-scale production due to its operational complexity. Here, we expressed and purified HA_t_ and S_t_ in both WT and *#3* plants. Western blot analysis showed comparable expression levels of the target proteins in WT and *#3* background, while HMC aggregates above the expected bands were consistently detected in WT but not in *#3* plants (Figure 3A, 3B, 3E and 3F). To assess purification efficiency in WT and *#3* plants, we expressed the S_t_ protein in both backgrounds, purified it from equal amounts of leaf tissue (1 g) using Ni²⁺-NTA resin, and analyzed the recovered proteins by SDS–PAGE followed by CBB staining. Quantification showed that the yield of S_t_ purified from *#3* was 70% higher than that from WT plants (Figure 3C and 3D). Using the same approach for HA_t_, we found a 40% increase in recovery from *#3* compared with WT (Figure 3G and 3H). Interestingly, the S_t_ or HA_t_ proteins purified from WT plants appeared as more diffuse bands with slightly higher apparent molecular weights on SDS–PAGE. We speculate that this anomaly is most likely caused by polyphenol binding (Figure 3C and 3G).

**Figure 2.**
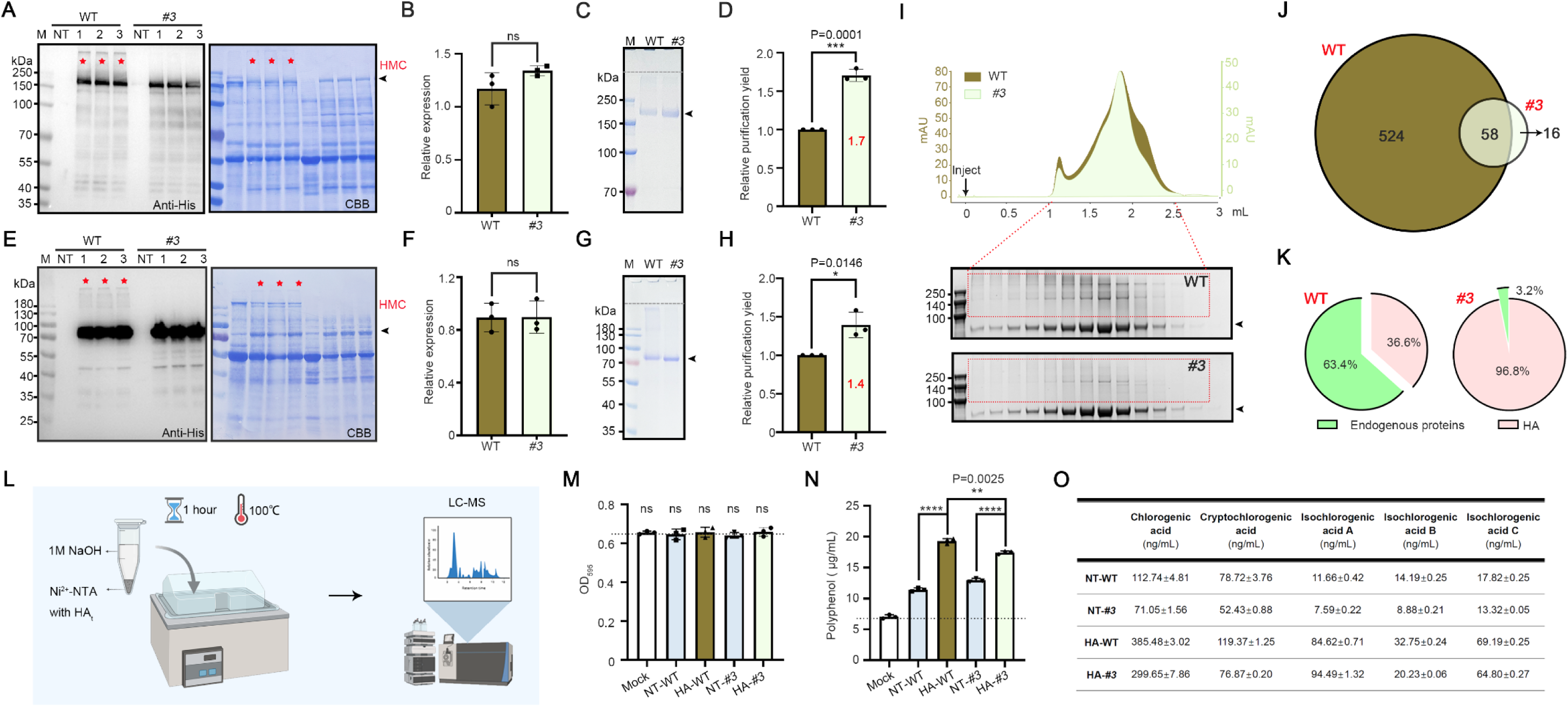
Purification of recombinant S_t_ and HA_t_ from *ppoa;ppob* plants shows increased purity and yield. **(A)** Western blot and CBB staining of S_t_ expressed in WT and *ppoa;ppob*#3 plants. NT, non-treated control. Red asterisks mark aggregated HMC. **(B)** Quantification of S_t_ expression in WT and *ppoa;ppob*#3 plants, based on band intensity in **(A)**. **(C)** CBB analysis of S_t_ purified from WT and *ppoa;ppob*#3 plants. Black arrows indicate the recombinant S_t_ bands. S_t_ was purified from 5 g of infiltrated leaf tissue of WT or *ppoa;ppob*#3 using Ni²⁺–NTA resin. One-hundredth of the purified S_t_ from each sample was analyzed by SDS–PAGE and CBB staining. **(D)** Relative purification yield of S_t_ in WT and *ppoa;ppob*#3 plants, based on band intensity in **(C)**. **(E)** Western blot and CBB staining of HA_t_ expressed in WT and *ppoa;ppob*#3 plants. NT, non-treated control. Red asterisks mark aggregated high molecular complexes. **(F)** Quantification of HA_t_ expression in WT and *ppoa;ppob*#3 plants, based on band intensity in **(E)**. **(G)** CBB analysis of HA_t_ purified from WT and *ppoa;ppob*#3 plants. Black arrows indicate the recombinant HA_t_ bands. HA_t_ was purified from 5 g of infiltrated leaf tissue of WT or *ppoa;ppob*#3 using Ni²⁺–NTA resin. One-hundredth of the purified HA_t_ from each sample was analyzed by SDS–PAGE and CBB staining. **(H)** Relative purification yield of HA_t_ in WT and *ppoa;ppob*#3 plants, based on band intensity in **(G)**. **(I)** Size exclusion chromatography (SEC) to the HA_t_ purified from WT and *ppoa;ppob*#3 plants, following the SDS-PAGE and CBB staining analysis to the fractions. Black arrows indicate the recombinant HA_t_ bands. Red dashed box indicates the high molecular complex (HMC); **(J)** LC-MS analysis to the HMC collected from WT and *ppoa;ppob*#3 after SEC and SDS-PAGE. Brown color indicates the proteins being detected from the HMC collected from WT; light green color indicates the proteins being detected from the HMC collected from *ppoa;ppob*#3. **(K)** Quantification of plant endogenous proteins and target protein (HA_t_) content of HMC collected from WT and *ppoa;ppob*#3 after SEC and SDS–PAGE. **(L)** Ni²⁺-NTA resin (50 μL) was incubated with total soluble extracts from 1 g of leaf tissue of WT and *ppoa;ppob*#3 plants with or without expressing HA_t_ for 15 min, followed by three times washing. Bound polyphenols were stripped by adding 400 μL of 1 M NaOH and heating at 100 °C. The supernatant was collected for subsequent Bradford assay **(M)** and polyphenol detection **(N)**. NT-WT, Non-infiltrated WT; HA-WT, HA_t_ infiltrated WT; NT-*#3*, Non-infiltrated *ppoa;ppob*#3; HA-*#3,* HA_t_ infiltrated *ppoa;ppob*#3. Dotted lines indicate the positive cutoff value, calculated as the mean titer of the Mock group. Statistical differences were determined by t-test. \*\**p* < 0.01; \*\*\*\**p* < 0.0001; ns, no significant. Error bars represent SEM. **(O)** Mass spectrometry detection of chlorogenic acid and its isomers in the four groups.

**Figure 3.**
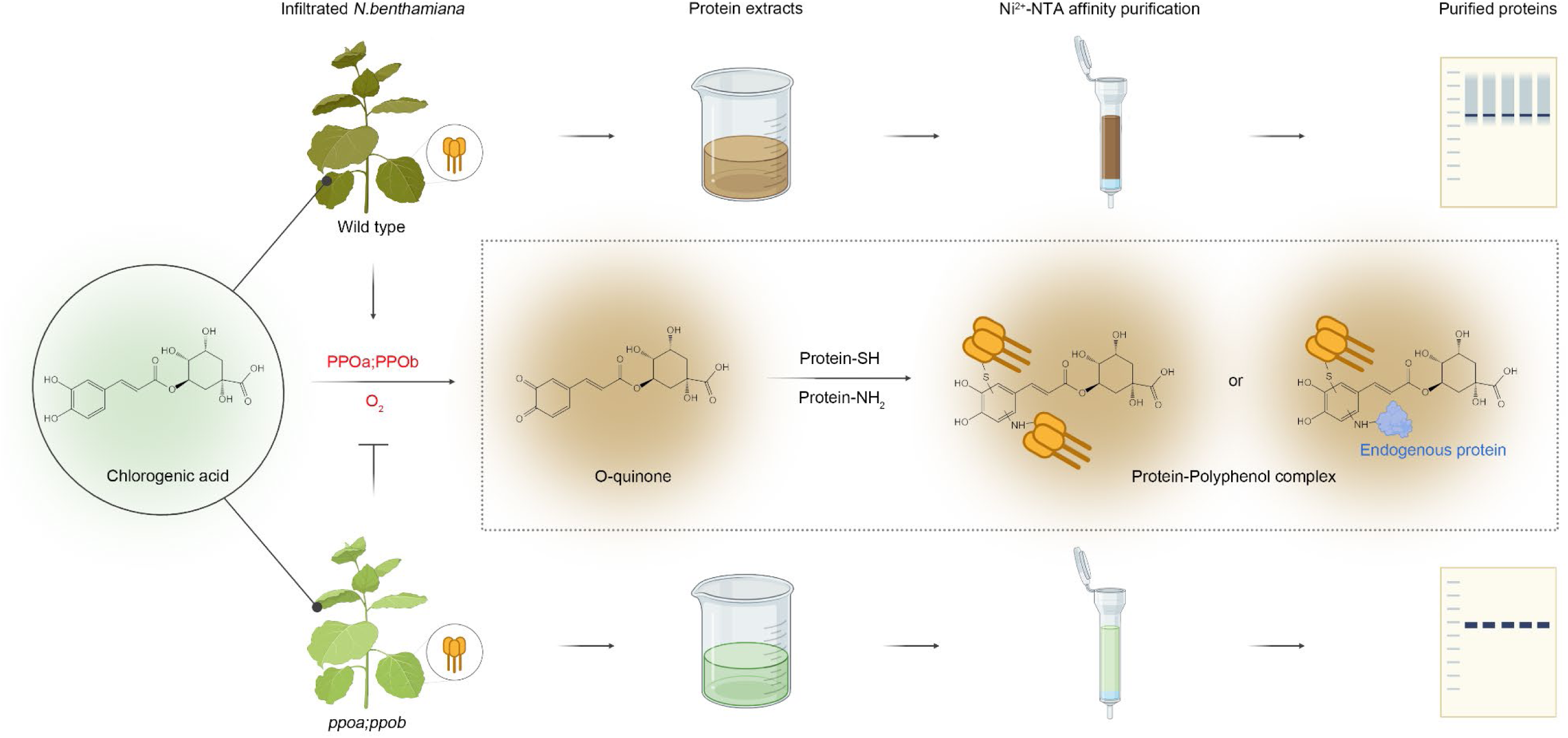
**Mechanism diagram of enzymatic browning and protein aggregation during recombinant protein purification process in *N. benthamiana*.**

To investigate the underlying cause of the increased purification yield, we analyzed both the fraction of total soluble protein that failed to bind to the affinity resin (unbound fraction) and the fraction that remained bound but could not be eluted from Ni^2+^-NTA beads. We found that HA_t_ proteins derived from the WT line exhibited substantially higher levels in both the unbound and non-eluted fractions, indicating that a significant portion of WT-derived proteins either did not interact efficiently with the resin or were irreversibly retained, whereas *#3*-derived proteins showed more efficient binding and recovery (Figure S2).

We next examined the quality of the purified HA_t_ proteins by size-exclusion chromatography, followed by SDS–PAGE of the eluted fractions. HA_t_ purified from WT plants contained markedly more HMC than that from *#3* (Figure 3I). Mass spectrometry analysis identified these aggregates as assemblies of the target HA_t_ protein with a range of endogenous plant proteins. In total, 581 distinct endogenous plant proteins were detected in the HMCs derived from WT plants, whereas only 73 distinct endogenous plant proteins were found in those from *#3* (Figure 3J). Protein quantification revealed that HMCs from WT contained 63.4% endogenous plant proteins and only 36.6% HA_t_, while HMCs from *#3* were composed almost entirely of the target protein, with 96.8% HA_t_ and only 3.2% endogenous contaminants (Figure 3K). Together, these results demonstrate that knocking out *PPOa* and *PPOb* greatly reduces host protein contamination and significantly improves both the yield and purity of recombinant proteins.

To examine whether polyphenols contribute to aggregate formation, we purified HA_t_ protein from both WT (HA-WT) and *#3* (HA-*#3*) plants using Ni²⁺-NTA resin. Non-infiltrated WT (NT-WT) and *#3* (NT-*#3*) plants were included as controls. After protein binding, the resin was washed and treated with 1 M NaOH and heated for one hour to strip polyphenols that were bound either to the recombinant protein or to the resin (Figure 3L). Bradford assays showed that all the NaOH eluates contained no detectable proteins, confirming that the treatment did not strip proteins (Figure 3M). In contrast, polyphenol detecting assays revealed a significantly higher amount of polyphenols in the eluates from HA-WT than from HA-*#3*(Figure 3N), which is identical with the HMC status. Low levels of polyphenols were also detected in NT-WT and NT-*#3* controls, suggesting that a small fraction of polyphenols may bind directly to the resin or to nonspecifically bound endogenous proteins. However, these background levels were significantly lower than the polyphenol content detected in HA-WT and NT-*#3* (Figure 3N).

Studies have reported that chlorogenic acid, rutin, and scopolamine together constitute more than 80% of the total polyphenol content, making them the predominant polyphenols in tobacco leaves(Zou et al., 2021). We next examined the residual levels of chlorogenic acid (Figure 3O) and other polyphenols (Figure S4) by mass spectrometry and detected multiple residual polyphenols across all four groups, including chlorogenic acid, catechin, benzoic acid, etc., indicating that protein–polyphenol interactions are non-selective. Notably, chlorogenic acid and its structural isomers were found to bind to recombinant proteins in the greatest amount, consistent with its predominance among polyphenols in tobacco. Taken together, we detected residual polyphenols in purified plant-derived recombinant proteins, with mass spectrometry identifying chlorogenic acid as the predominant species. The residual levels correlated positively with HMC content, further confirming that plant-derived recombinant proteins are subject to polyphenol binding and contamination during purification.

## Discussion

Unlike microbial or mammalian platforms, plants such as *Nicotiana benthamiana* produce a complex array of secondary metabolites, particularly polyphenols, which interfere with protein extraction during downstream processing. Our study demonstrates that oxidation of these metabolites by PPOs is a major factor driving protein aggregation during purification, thereby compromising both protein quality and yield. By genetically eliminating PPO activity, we show a clear reduction in protein–polyphenol crosslinking, which improves the integrity and recoverability of recombinant proteins (Figure 3). These results provide direct genetic and biochemical evidence linking PPO-mediated oxidation to the challenges of plant-based protein purification. Importantly, our work establishes a cleaner and more scalable *N. benthamiana* chassis, addressing a long-standing barrier in plant molecular farming.

By combining transcriptomic profiling with VIGS, we identified *PPOa* and *PPOb* as the primary enzymes responsible for polyphenol oxidation in *N. benthamiana*. Silencing both genes was necessary to suppress enzymatic browning and protein aggregation, pointing to functional redundancy. Building on this insight, we generated *ppoa;ppob* double knockout mutants and validated their performance across multiple generations. Although early growth delays were observed, mature plants showed no significant developmental defects, indicating that the knockout lines are viable and suitable for production-scale applications.

Our Western blot analysis comparing the expression levels of GFP, HA_t_, and S_t_ in WT and *#3* lines revealed no substantial differences. This observation contrasts with previous reports, in which GFP fluorescence quantification demonstrated a significant increase in GFP expression in PPO-silenced lines(Mahadevan et al., 2025). We suspect that this discrepancy may arise from the experimental system: in our hands, VIGS caused highly variable effects on plant growth, which could in turn influence the accumulation of heterologous proteins.

Crucially, while expression levels in wild-type and mutant plants were comparable, proteins purified from the mutant lines exhibited higher recover efficiency and homogeneity. However, even in the double-knockout background, trace amounts of HMC could still be detected on Coomassie-stained SDS-PAGE gels. Smaller amounts of polyphenol were also detectable using polyphenol assay kits and mass spectrometry. This residual presence may be attributed to two factors: 1) polyphenols that were not oxidized by PPOs may bind to the proteins or Ni^2+^-NTA resin through noncovalent interactions rather than covalent crosslinking; 2) PPOa and PPOb, while the major contributors, are not the only polyphenol oxidases carrying out this function. To further reduce polyphenol contamination, purification conditions could be optimized to disrupt noncovalent interactions. Additionally, further knockouts of *PPOc* and *PPOd* on the *ppoa;ppob* background could be performed to assess whether polyphenol contamination can be completely eliminated.

## Methods

### Construction of recombinant genes

DNA fragments encoding GFP/HA/Spike together with the trimerization motifs of the mouse Coronin 1A (mCor1; VSRLEEDVRNLNAIVQKLQERLDRLEETVQAK) were C-terminal fused with the His-tag and the ER retention signal HDEL. The ER leader peptide of Arabidopsis BIP1 was N-terminally fused with all the constructs for ER targeting. The GFP gene(Song et al., 2022a), HA gene of swine influenza virus(Song et al., 2025) and Spike gene of SARS-Cov-2(Song et al., 2022b) were sub-cloned from our published plasmids via *BamHI* and *XmaI*. All the sequences used in these studies were attached to Table 1.

**Table 1.**
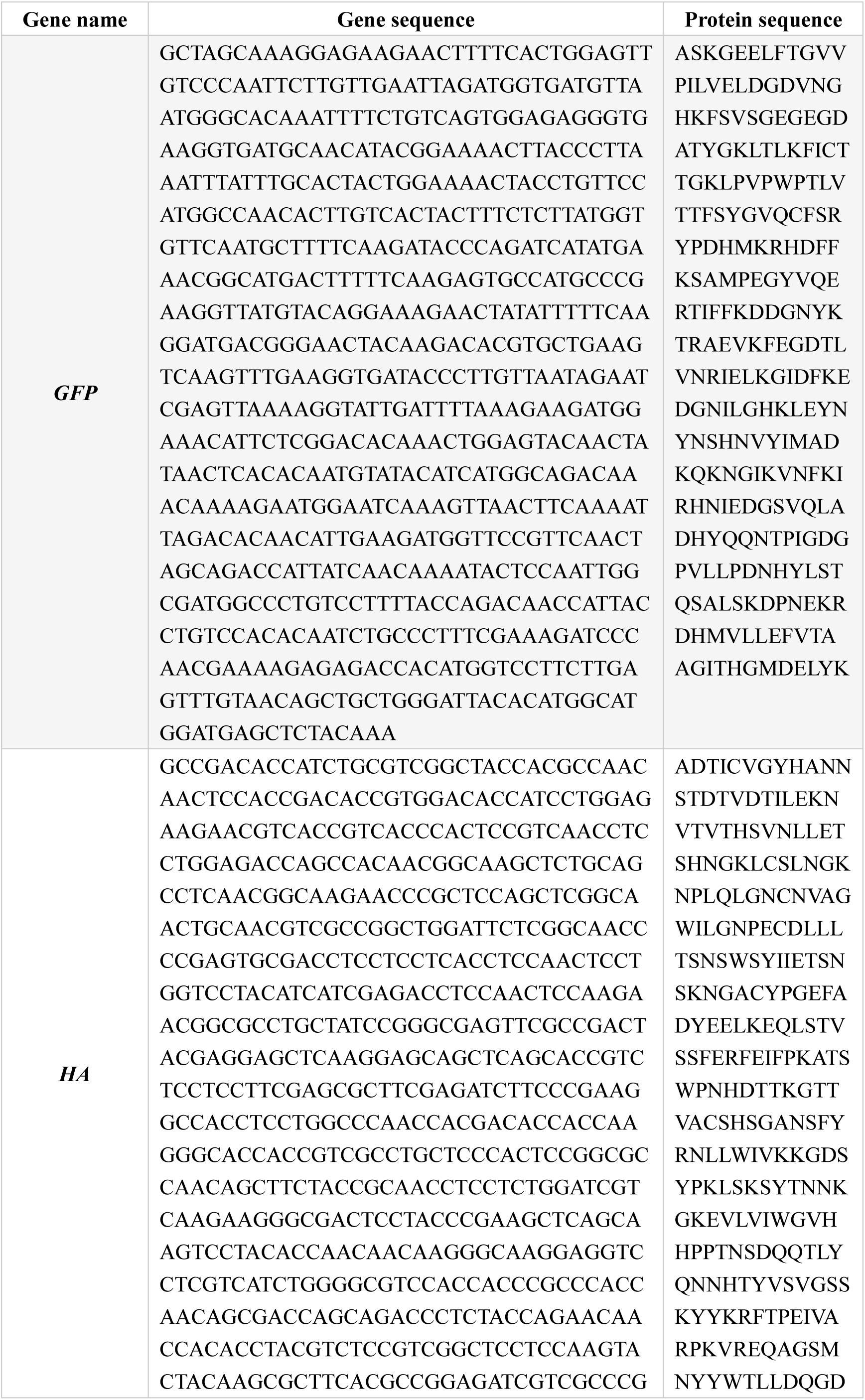
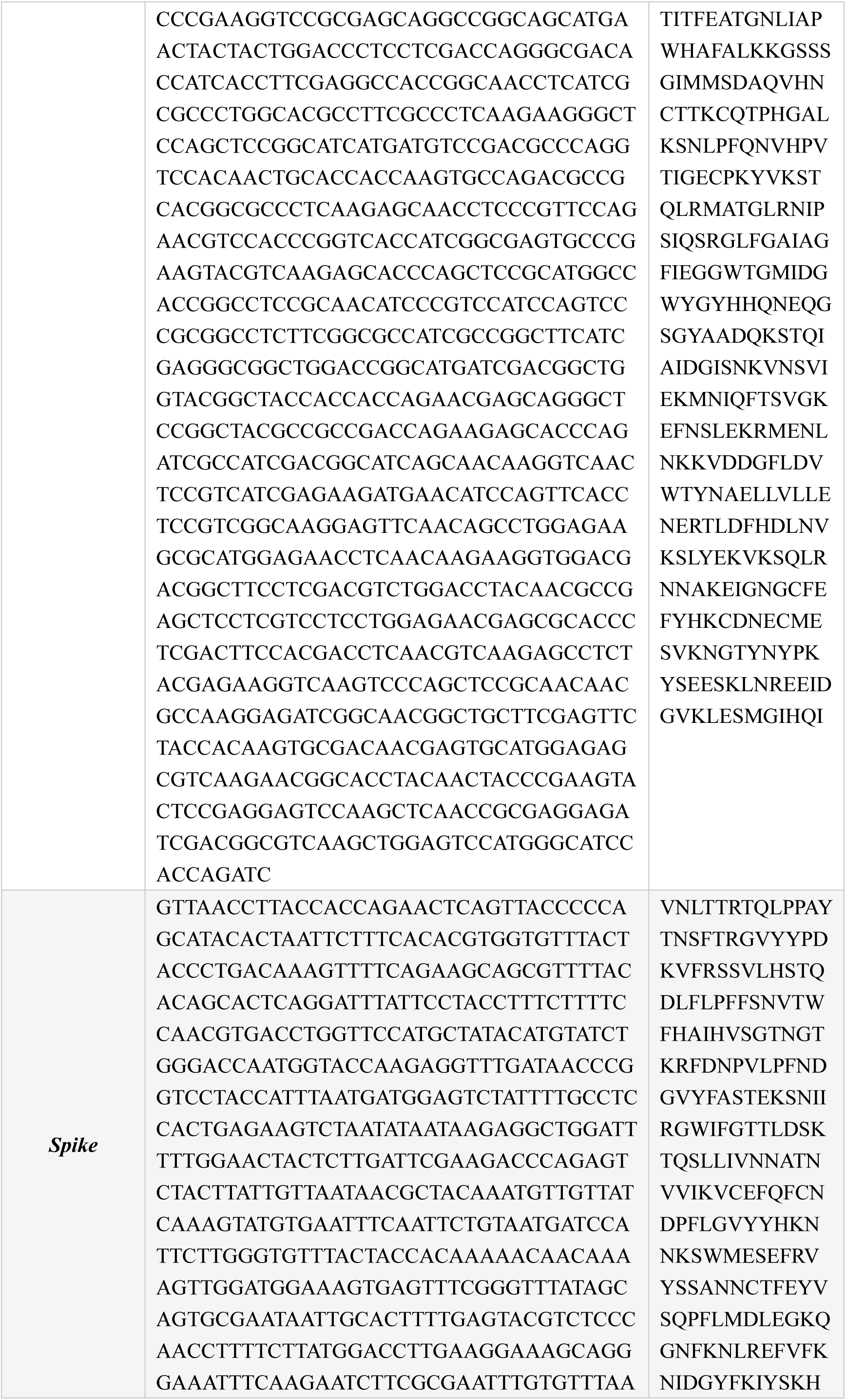
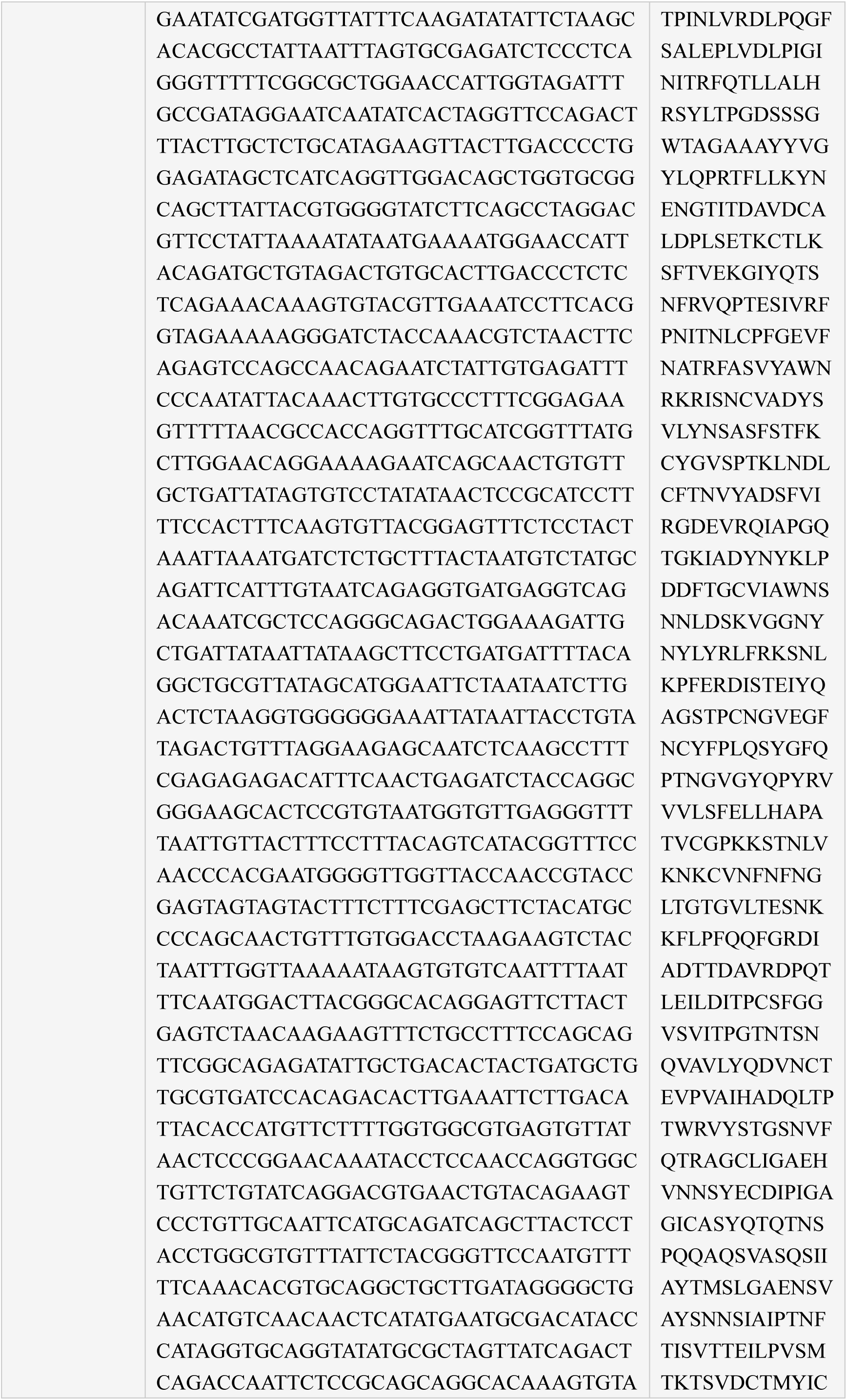
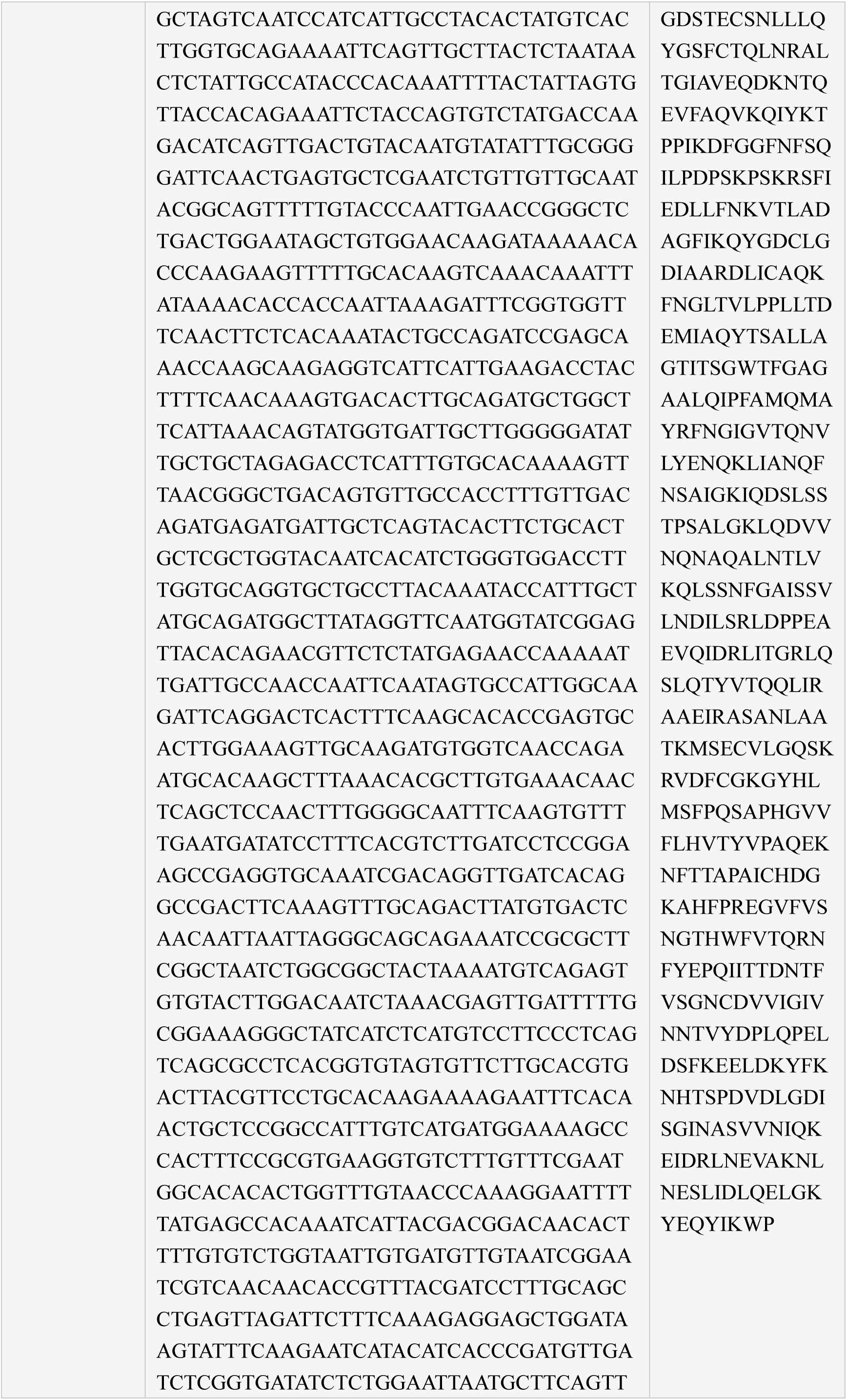
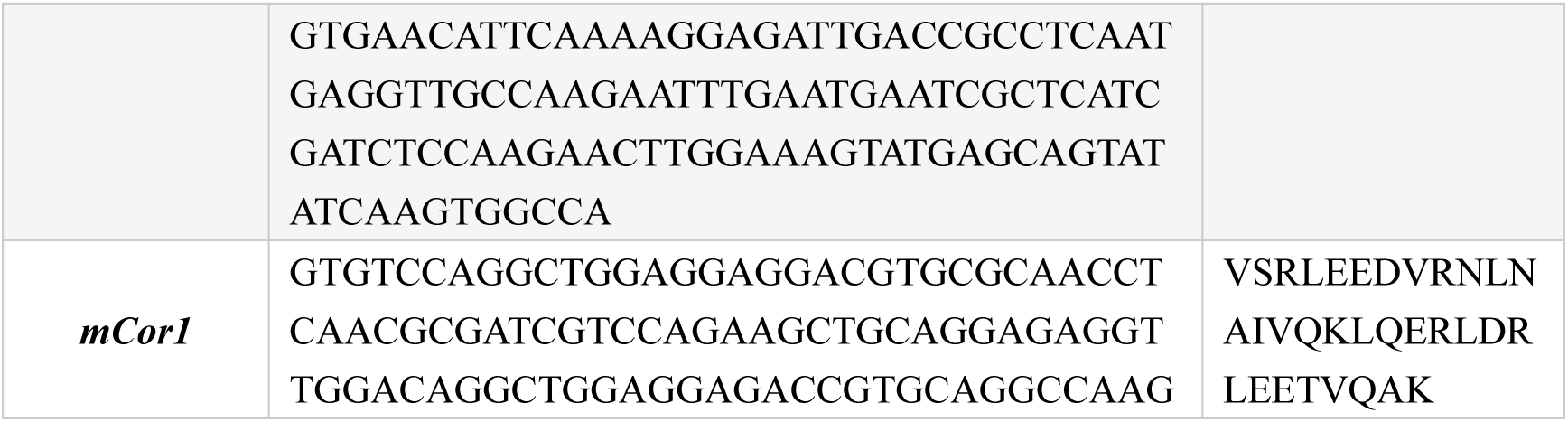
Gene and protein sequences used in this study.

### Production of transient transgenic plants

Expression vectors were introduced into *Agrobacterium* strain GV3101 by electroporation. A single colony of *Agrobacterium* harbouring expression vectors was inoculated to YEP Broth and cultured in an incubator at 28℃ overnight. 4-week-old *N. benthamiana* plants grown in a greenhouse at 25°C with a 16 h light/8 h dark cycle were used for Agroinfiltration by syringe. The infiltrated leaves were harvested at three-days post infiltration (DPI) to examine the expression level. A vacuum chamber is used for the large-scale production of Agroinfiltration.

### Virus-induced Gene Silencing (VIGS)

Agrobacterium containing TRV1 or TRV2 were grown overnight at 28°C in YEP medium containing 25 mg/L rifampicin and 50 mg/L kanamycin. The cultures were centrifuged at 5000 x *g* for 10 minutes at room temperature and the pellet was resuspended at OD_600_ = 1.0 in agroinfiltration buffer (10 mM MES pH 5.7, 10 mM MgCl_2_, 100 µM acetosyringone). TRV2 cultures were mixed at a 1:1 ratio with TRV1 cultures. Two-week-old *N. benthamiana* were agroinfiltrated with the bacterial suspension mixture using a 1 ml needleless syringe. Three to five weeks later plants were assessed for silence by observing whether the *TRV2::PDS* plants had bleached leaves. Silenced plants were then tested by agroinfiltration. Primers used for VIGS were listed in Table 2.

**Table 2.**
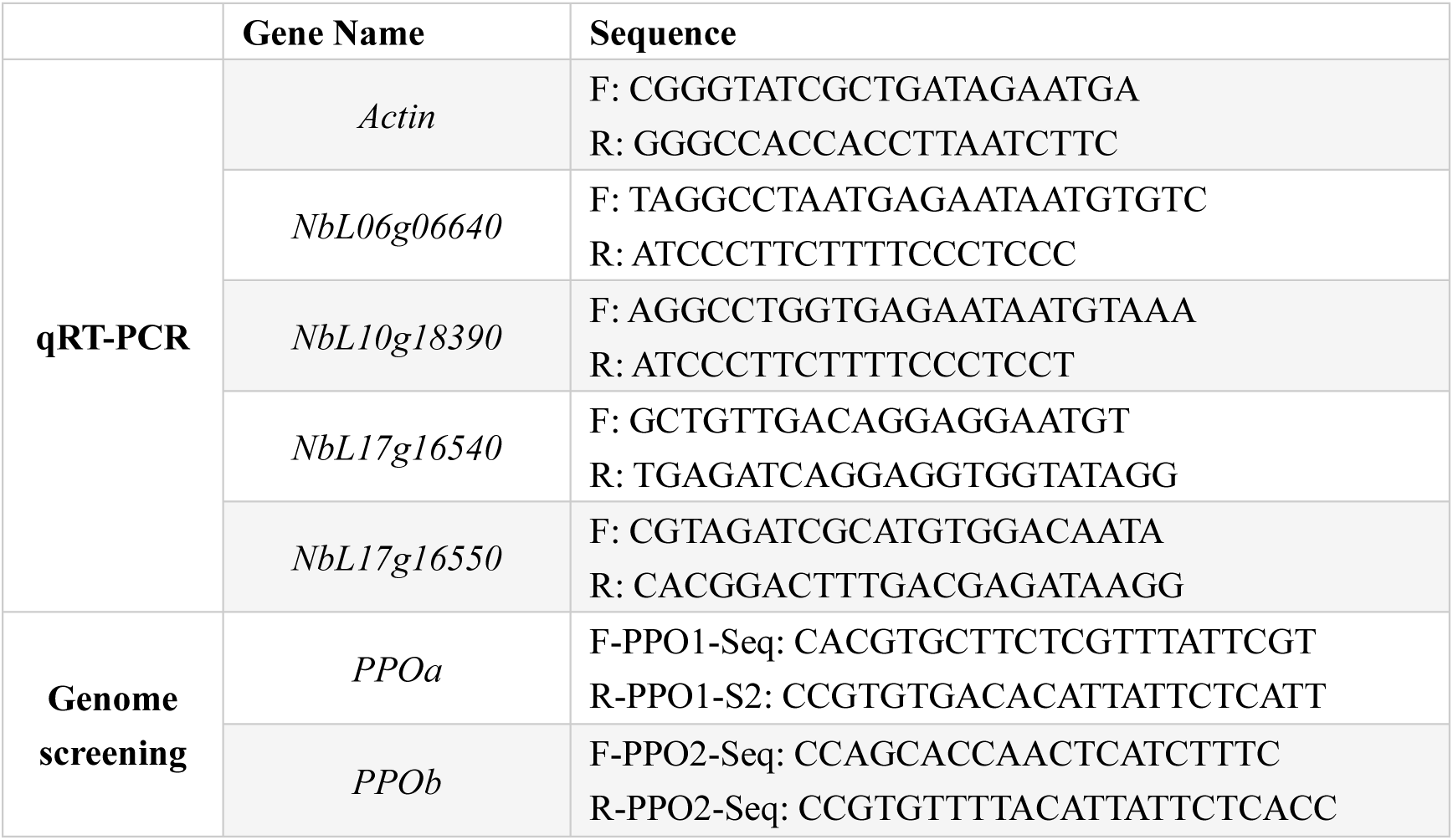
Primers used in this study.

**Table 3.** Detection of additional polyphenols after purification.

|  | Rutin<br>(ng/mL) | Protocatechualdehyde<br>(ng/mL) | Catechin<br>(ng/mL) | Epicatechin<br>(ng/mL) | Caffeic acid<br>(ng/mL) | Vanillin<br>(ng/mL) | Quercetin 3-β-D-glucoside<br>(ng/mL) | Benzoic acid<br>(ng/mL) | Quercetin<br>(ng/mL) |
| --- | --- | --- | --- | --- | --- | --- | --- | --- | --- |
| **NT-WT** | 0.45±0.04 | 12.55±0.58 | 9.57±0.27 | 13.79±0.19 | 2.29±0.15 | 6.36±0.22 | 2.04±0.15 | 119.94±1.83 | 1.95±0.06 |
| **NT-#3** | 3.045±0.27 | 9.97±0.59 | 11.78±0.31 | 17.35±0.12 | 2.79±0.07 | 3.77±0.29 | 9.65±0.06 | 65.17±2.5 | 6.94±0.35 |
| **HA-WT** | 3.11±0.15 | 41.94±0.82 | 100.99±2.07 | 35.69±1.65 | 16.80±0.15 | 12.07±0.09 | 14.90±0.43 | 44.79±2.22 | 3.21±0.08 |
| **HA-#3** | 1.99±0.18 | 55.63±2.12 | 84.82±4.53 | 30.63±1.59 | 12.35±0.18 | 6.24±0.13 | 9.61±0.16 | 48.25±1.51 | 1.66±0.06 |

### RNA extraction and qRT-PCR

Total RNA was extracted from 4-week-old *N. benthamiana* leaf tissue using RNeasy Plant Mini Kit (Qiagen, Cat no.74904). Approximately 100 mg of fresh leaf tissue was ground to a fine powder in liquid nitrogen for this experiment following the specification. RNA quantity and purity were assessed by spectrophotometry, and integrity was confirmed by agarose gel electrophoresis. First-strand cDNA synthesis was carried out using the Hifair® Ⅲ 1^st^ Strand cDNA Synthesis SuperMix for qPCR kit (Cat#11141ES60). The qRT-PCR quantification experiment was performed using reagent kits from YEASEN Biotechnology (Shanghai) Co., Ltd. Fluorescence-based quantitative PCR detection was conducted using the Hieff® qPCR SYBR Green Master Mix kit. Primers used for qRT-PCR were listed in Table 2.

### Genome editing of *PPO* and screening mutants

The *PPOa* and *PPOb* knockout lines were constructed using the CRISPR-Cas9 genome editing system. Two gRNAs were designed to target the homologous sequences of *PPOa* and *PPOb,* which is ACCTTGACGCTGTTGACAGGAGG and GTTAGCCACTAGTCGAATGAGGG respectively. Both target sites were in the N-terminal coding regions. Following *Agrobacterium-*mediated transformation and tissue culture, a total of 52 transgenic lines were obtained. These transgenic seedlings were transferred from culture medium to soil for further growth. Genomic DNA was extracted from the leaves of T1 generation plants and subjected to sequencing analysis. Primers used for PCR and sequencing are listed in Supplementary Table 2. Among the 52 lines, two double knockout mutants (*ppoa;ppob*), designated as lines *#3* and *#7*, were identified. In addition, three single knockout mutants of *PPOa* were also isolated.

### SDS-PAGE and western blot analysis

Infiltrated leaves were ground and homogenized in protein extraction buffer (TBS buffer containing 1 mΜ EDTA, 0.2% Tween 20(v/v), and 1mM PMSF). The total protein extracts or purified proteins were separated by 8% or 10% SDS-PAGE. Western blot analysis was performed using the mouse anti-His antibody (1: 10000 dilutions, CWBIO, Cat#CW0286M). The secondary antibodies used in this study were sheep anti-Mouse IgG conjugated HRP (1: 5000 dilution, Abcam, Cat#ab6789) Immunoblots were developed with the enhanced chemiluminescence kit (TantonTM High-sig ECL Western Blotting Substrate, Cat#180-501), and images were captured using the Tanon 5200 Multi system (Shanghai, China).

### In-gel digestion of HMC

To identify the proteins in HMC, high-resolution liquid chromatography–tandem mass spectrometry (LC-MS/MS) analysis was performed following in-gel digestion. Protein bands of interest were excised and destained with 50% acetonitrile in 50 mM ammonium bicarbonate, dehydrated with acetonitrile, and subjected to reduction with 10 mM DTT at 56 °C for 1 h and alkylation with 20 mM iodoacetamide at room temperature in the dark. After dehydration and drying, in-gel digestion was carried out overnight at 37 °C using trypsin (0.025 μg/μL), followed by peptide extraction with 5% TFA, 50% acetonitrile, and 45% water.

### Tandem MS Analysis for HMC

The LC-MS/MS was performed for this study using an Orbitrap Fusion Lumos instrument (ThermoFisher) coupled with an Vanquish Neo nano HPLC (ThermoFisher). Mobile phase A and B were water and 80% acetonitrile, respectively, with 0.1% formic acid. Protein digests were loaded directly onto a self-packed to a 17 cm C18 column with 1.9 μm Reprosil PUR beads (Dr.Maisch HPLC GmbH) running at a flow rate of 600 nL/min. Digested peptides were separated at 600 nL/min using the following gradient: 0-2 min, 4-8%B; 2-35min, 8%-28%B; 35-55 min, 28-40%B; 55-56 min, 40-95%B; 56-66 min, 95%B; Data were acquired in data-dependent MS/MS mode.

Mass spectrometry data were acquired in data-dependent acquisition (DDA) mode on a Q Exactive mass spectrometer. The full MS (MS¹) parameters were as follows: resolution, 70,000; AGC target, 3 × 10⁶; maximum injection time (IT), 100 ms; and scan range, m/z 300–1800. For tandem MS (MS²), the parameters were: resolution, 17,500; AGC target, 1 × 10⁵; maximum IT, 50 ms; TopN, 20; and normalized collision energy (NCE, stepped), 28. Raw spectral data were generated after acquisition.

Database searching was performed with MaxQuant against the *Nicotiana benthamiana* (*NbLab360.v103*) database(Ranawaka et al., 2023). The search parameters were set as follows: fixed modification, carbamidomethylation (C); variable modifications, methionine oxidation (M) and protein N-terminal acetylation; enzyme specificity, trypsin; maximum missed cleavages, 2; peptide mass tolerance, 20 ppm; and fragment mass tolerance, 20 ppm.

### Ni^2+^-NTA binding assay

We perform this experiment mentioned in our previous study(Song et al., 2021). Briefly, the plant tissues were well ground in liquid nitrogen and dissolved in extraction buffer (Tris 50 mM, pH 7.5, NaCl 150 mM, imidazole 5 mΜ, 0.1% Tween 20, and PMSF 1mM, v/m = 10: 1), followed by centrifugation two to four times to remove the debris as much as possible. The supernatant filtered by 0.22μM bacterium-free filter unit were incubated with Ni^2+^-NTA beads at 4°C for 15 mins. Then, the beads were collected after washing with washing buffer (Tris 50 mM, pH 7.5, NaCl 150 mM, imidazole 10 mΜ, 0.2% Tween 20). Finally, the conjugated proteins were eluted using elution buffer (Tris 50 mM, pH 7.5, NaCl 150 mM, imidazole 200 mΜ, 0.2% Tween 20), followed by imidazole dialysis using 30 kDa cutoff Laboratory Concentration Devices (Merck, UFC9030).

### Polyphenol stripping and detection

Approximately 1 g of leaf tissue from *N. benthamiana* (non-transformed and HA-infiltrated) and *ppoa;ppob#3* plants (non-transformed and HA infiltrated) was homogenized and subjected to Ni²⁺-NTA binding according to the procedure described in the “Ni²⁺-NTA Binding Assay” section of this paper. After binding, the beads were washed sequentially with washing buffer (50 mM Tris-HCl, pH 7.5; 150 mM NaCl; 10 mM imidazole; 0.1% Tween-20) and deionized water. Residual liquid was carefully removed. Subsequently, 500 µL of 1 M NaOH was added to the Ni²⁺-NTA beads, followed by incubation at 100 °C for 1 hour. The supernatant was collected by centrifugation. Polyphenol content in the resulting supernatant was quantified using the Plant Total Phenol Content Assay Kit (Solarbio, Cat# BC1340). Protein concentration was determined in parallel using the Ready-to-use Bradford Protein Assay Kit (Detergent Compatible; EpiZyme, Cat# ZJ104).

### Polyphenol samples extraction for LC/MS

Samples were extracted with 0.5 mL of 80% methanol containing 0.2% vitamin C using ultrasonic treatment for 30 min, followed by centrifugation at 12,000 rpm for 10 min. The supernatant was collected, and the extraction was repeated three times, after which all supernatants were pooled. The extracts were analyzed using a UPLC-Orbitrap-MS system (UPLC, Vanquish; MS, Q Exactive, Thermo Fisher Scientific). Chromatographic separation was performed on a Waters HSS T3 column (50 × 2.1 mm, 1.8 μm) at 40 °C, with a flow rate of 0.3 mL/min and an injection volume of 2 μL.

### UPLC conditions and LC-MS

For detection of chlorogenic acid and its isoforms, the samples were analyzed using an UPLC-Orbitrap-MS system (UPLC, Vanquish; MS, QE). The analytical conditions were as follows, UPLC: column, Waters ACQUITY UPLC HSS T3(1.8 μm, 2.1 mm×50 mm); column temperature, 40 ℃; flow rate, 0.3 mL/min; injection volume, 2 μL; solvent system, water (0.1% acetic acid): acetonitrile (0.1% acetic acid); gradient program, 97:3 V/V at 0 min, 97:3 V/V at 1.0 min, 50:50 V/V at 5.0 min, 10:90 V/V at 6.0 min, 10:90 V/V at 7.0 min, 97:3 V/V at 7.1 min, 97:3 V/V at 9.0 min. HRMS data were recorded on a Q Exactive hybrid Q-Orbitrap mass spectrometer equipped with a heated ESI source (Thermo Fisher Scientific) utilizing the SIM MS acquisition methods. The scan range for the primary MS was set to m/z 300–600(Jaiswal et al., 2014; Glauser et al., 2016).

For detection of polyphenols except chlorogenic acid and its isoforms, the sample extracts were analyzed using an UPLC-Orbitrap-MS system (UPLC, Vanquish; MS, QE). The analytical conditions were as follows, UPLC: column, Waters HSS T3 (50×2.1 mm, 1.8 μm); column temperature, 40℃; flow rate, 0.3 mL/min; injection volume, 2 μL; solvent system, water (0.1% formic acid): acetonitrile (0.1% formic acid); gradient program, 90:10 V/V at 0 min, 90:10 V/V at 2.0 min, 40:60 V/V at 6.0 min, 40:60 V/V at 9.0 min, 90:10 V/V at 9.1 min, 90:10 V/V at 12.0 min. HRMS data were recorded on a Q Exactive hybrid Q-Orbitrap mass spectrometer equipped with a heated ESI source (Thermo Fisher Scientific) utilizing the Fullms-ms2 acquisition methods. The scan range for the primary MS was set to m/z 100–900(Glauser et al., 2016; Zhang et al., 2019).

## Author contributions

SJS contributed to the conception of the study and wrote the manuscript; IHH critically reviewed and revised the text. HPD and HXM contributed significantly to experiments and data analysis with the help of XJX. YFG offered good suggestions.

## Acknowledgements

This research was supported by the Science and Technology Foundation of Beijing Life Science Academy (2024200CB0110), the Natural Science Foundation of Shandong Province (ZR2024QC068).

---

**Figure S1.**
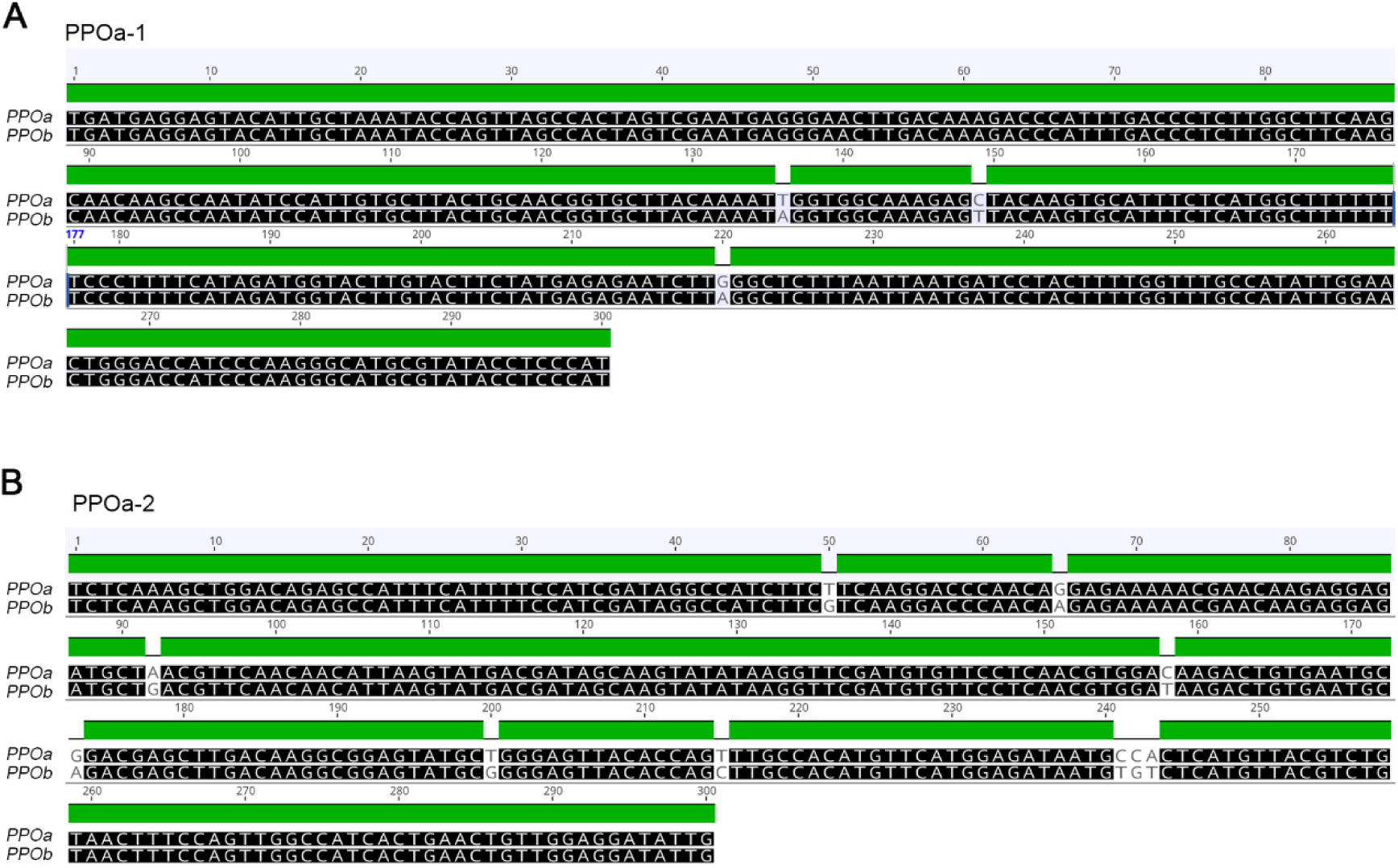
Sequence used for VIGS assay. **(A)** Sequence of PPOa-1(300nt) used for VIGS assay and its alignment to *PPOb*. **(B)** Sequence of PPOa-2(300nt) used for VIGS assay and its alignment to *PPOb*.

**Figure S2.**
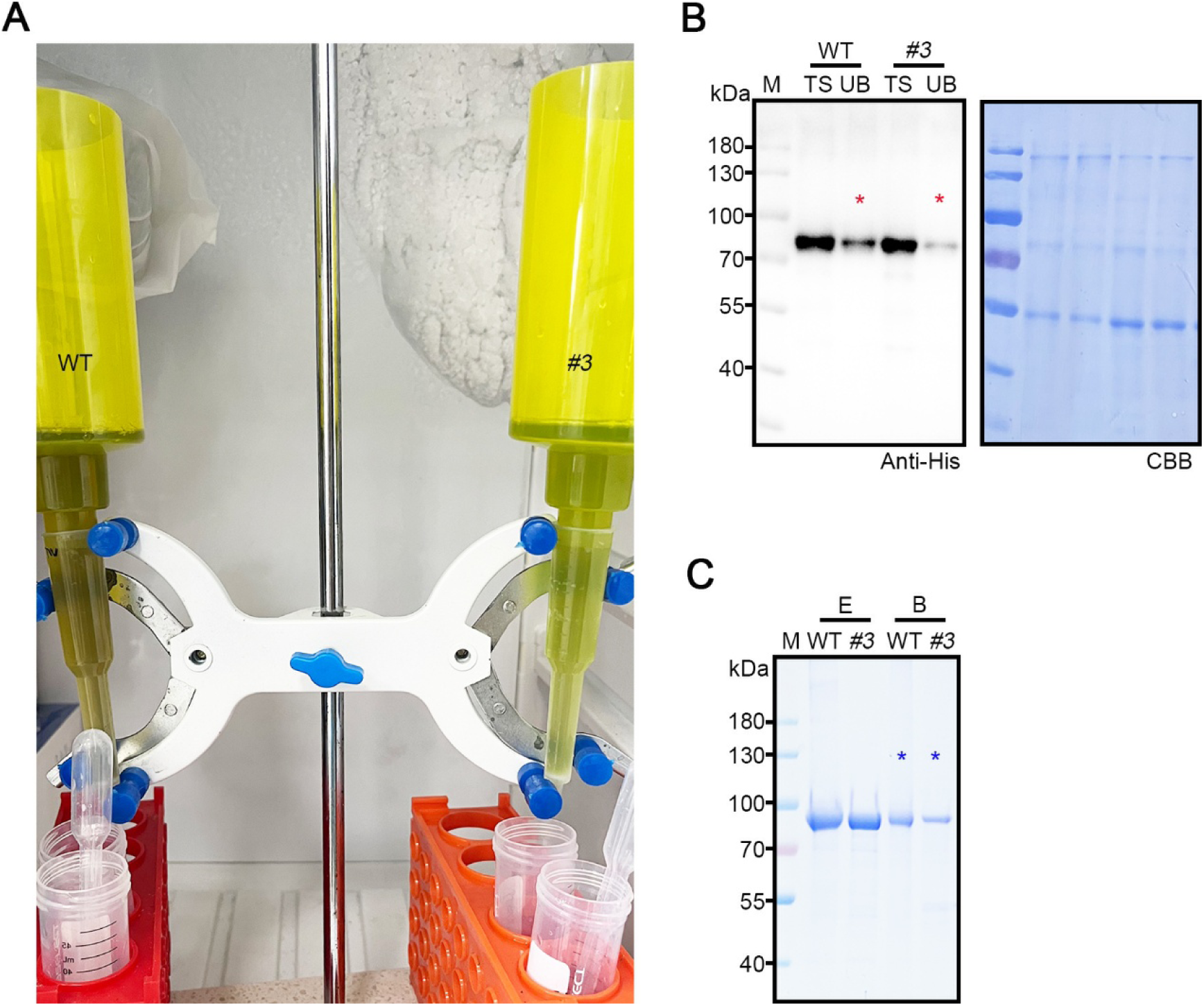
Detection of target proteins during Ni^2+^-NTA purification. **(A)** Schematic diagram of the gravity-flow Ni^2+^-NTA column purification setup. **(B)** Binding efficiency of HA_t_ proteins from WT and line *#3* to the Ni^2+^-NTA resin. TS, total soluble protein; UB, unbound protein. Red asterisks mark the unbound target proteins. **(C)** Elution efficiency of HA_t_ proteins from WT and line *#3* from the Ni^2+^-NTA resin. Blue asterisks mark the non-eluted proteins on the beads. E, elusion; B, beads.

**Figure S3.**
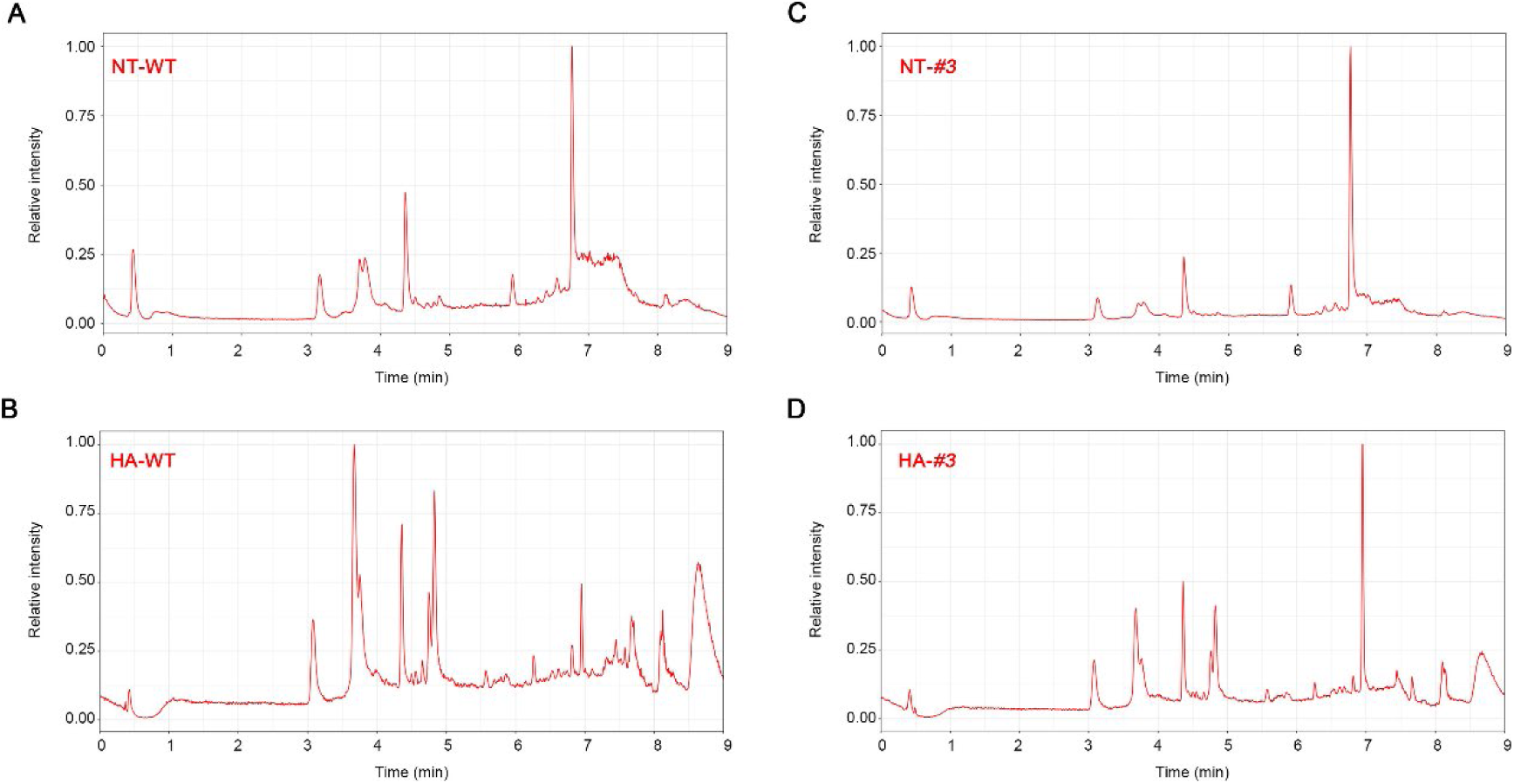
Total ion chromatogram (TIC) for the detection of chlorogenic acid and its isomers. Four samples (**A**, NT-WT; **B**, HA-WT; **C**, NT-*#3*; **D**, HA-*#3*) were analyzed using a UPLC-Orbitrap-MS system, respectively. UPLC conditions were shown as described in M&M. HRMS data were recorded on a Q Exactive hybrid Q-Orbitrap mass spectrometer equipped with a heated ESI source (Thermo Fisher Scientific) utilizing the SIM MS acquisition methods. The scan range for the primary MS was set to m/z 300–600.

**Figure S4.**
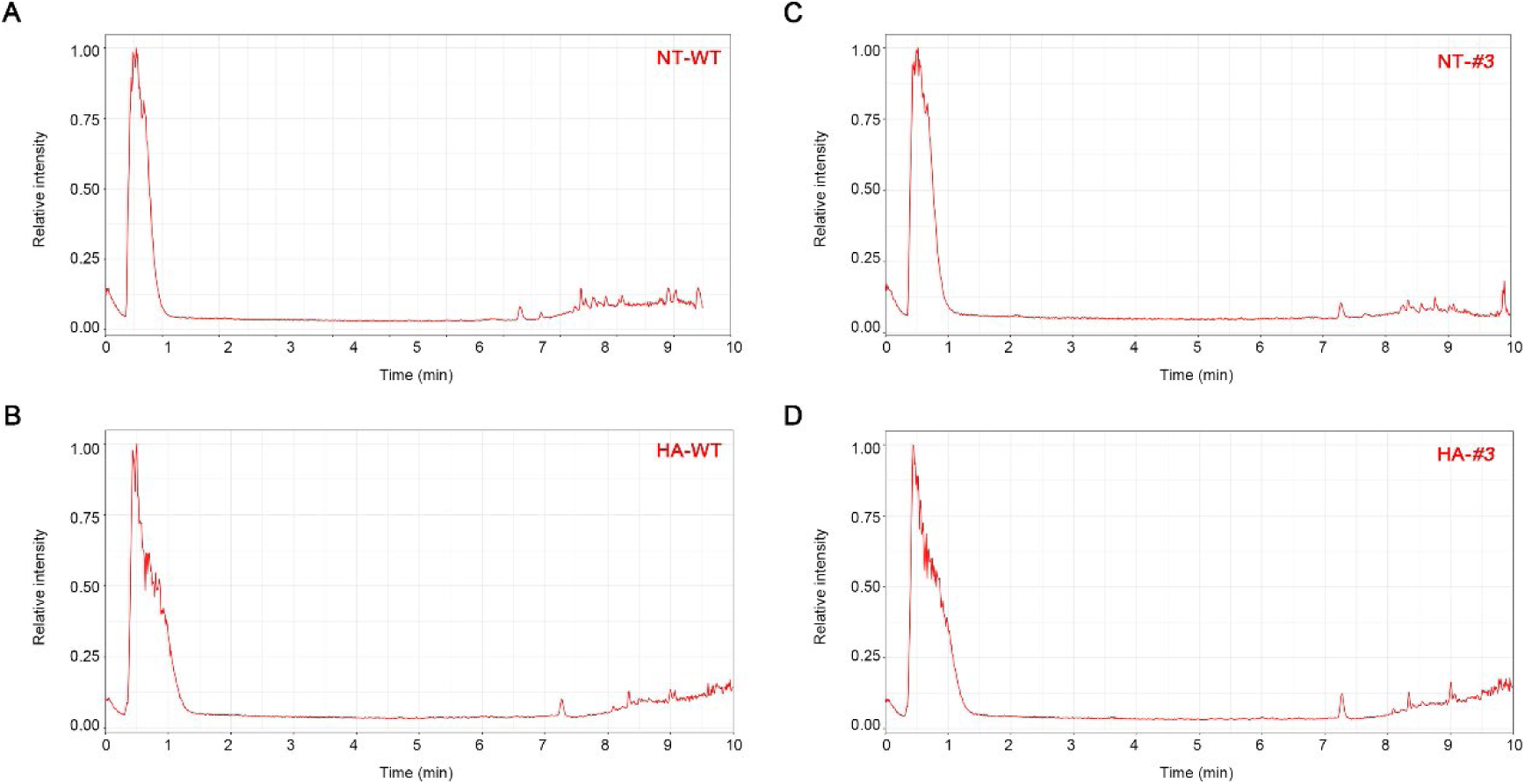
Total ion chromatogram (TIC) for the detection of other polyphenols except chlorogenic acid and its isomers. Four samples (**A**, NT-WT; **B**, HA-WT; **C**, NT-*#3*; **D**, HA-*#3*) were analyzed on a UPLC-Orbitrap-MS system. UPLC conditions were shown as described in M&M. HRMS data were recorded on a Q Exactive hybrid Q-Orbitrap mass spectrometer equipped with a heated ESI source (Thermo Fisher Scientific) utilizing the Fullms-ms2 acquisition methods. The scan range for the primary MS was set to m/z 100–900.

